# ConsensuSV-ONT - a modern method for accurate structural variant calling

**DOI:** 10.1101/2024.07.26.605267

**Authors:** Antoni Pietryga, Mateusz Chiliński, Sachin Gadakh, Dariusz Plewczynski

## Abstract

The improvements in sequencing technology make the development of new tools for the detection of structural variance more and more common. However, the tools available for the long-read Oxford Nanopore sequencing are limited, and it is hard to choose one, which is the best. That is why there is a need to create a tool based on consensus that combines existing work in order to discover a set of high-quality, reliable structural variants that can be used for further downstream analysis. The field has also been subject to revolution in machine learning techniques, especially deep learning. In the spirit of the aforementioned need and developments, we propose a novel, fully automated ConsensuSV-ONT algorithm. The method uses six independent, state-of-the-art structural variant callers for long-read sequencing along with a convolutional neural network for filtering high-quality variants. We provide a runtime environment in the form of a docker image, wrapping a nextflow pipeline for efficient processing using parallel computing. The solution is complete in its form and is ready to use not only by computer scientists but accessible and easy to use for everyone working with Oxford Nanopore long-read sequencing data.

## Introduction

The study of human variation has been an important topic in the research community since the first genome was assembled ^1^. With the development of new experimental methods, multiple tries have been done to analyse genomes on the population scale - including the 1000 Genomes project ^2^. New technologies allowed more accurate detection of the longest mutations - Structural Variants (SVs), which also pushed the community to propose new algorithms. The importance of the problem has been proven by many experimental studies, which show that the accurate detection of mutations such as deletions, insertions, inversions, translocations and other long variants can explain the causes of many diseases ^3,4^.

One of the most recent analyses based on a combination of PacBio, Bionano, and Illumina sequencing techniques pushed the detection possibilities to an average of 27,622 SVs per person ^5^, in comparison to 4405 reported by 1000 Genomes project.

The scientific community, for the accurate detection of variants, usually uses Illumina and PacBio sequencing technology ^6^ - while the first is used mostly for identifying short indels, and single nucleotide polymorphisms, the latter is much more suitable for the detection of long variants. However, there has been an increase in interest in Oxford Nanopore (ONT) sequencing technology ^7^, ^8^, as an alternative to the PacBio. As the method is being strongly developed, we decided to investigate the possibilities of increasing the accuracy and precision of long variant detection using ONT technology.

In both short- and long-read technologies many tools have been proposed for detecting structural variants. However, there is no universal tool that beats all the other ones - with every algorithm, we unfortunately have a bias, or method-dependent issues, that make us use multiple algorithms for the detection of SVs. That is why it has been shown that the use of consensus - combining the results from many SV callers - allows achieving better results than using individual tools - and is used both with Illumina ^9^ and PacBio sequencing ^10,11^. However, we have not found any tools that have been tested in detail for ONT sequencing technology. Additionally, the tools proposed for PacBio require the independent preparation of batch files. That is why we have decided to propose a unified calling platform by providing a full runtime environment (docker) and an entire pipeline that goes from generating alignment files through generating lists of variants using various SV callers to final filtering and scoring.

In our work, we present a novel meta-caller algorithm, along with a fully automated variant detection pipeline and a high-quality variant filtering algorithm based on variant encoding for images and convolutional neural network models. We tested the algorithm, called ConsensuSV-ONT, and other individual callers on three real datasets (HG00733, HG00514, NA19240).

## Results

### ConsensuSV-ONT overview

The algorithm we propose in this work is used as a method for detecting high-quality variants based on alignment. Consensus-ONT consists of four phases. First, a set of SV candidates is generated - a set of unique (non-overlapping) variants. The SV candidates collection is created based on existing tools and is automated and called within the pipeline. Using the Truvari ^12^ tool (merge module), we combine the result files from individual tools and remove overlapping variants using the collapse module. In order to train the model, we divided the combined results into two datasets, depending if the variant is also present in the ground truth set. Such division was done using the bench module of the Truvari tool. In the third step, each variant is encoded as an image (see Methods). To test the model, we used 3 ONT samples, and we took one as our training set, one as the validation set, and one as the final testing of the model. The data in the testing set is stored independently and is not used either for training or validating and choosing the model, ensuring the highest quality of the results. The final layer in the model is sigmoid, thus, the results of the model are the probability of the variant being detected. In our study, we have decided to choose 0.5 as the cut-off threshold for the variants to be considered as belonging to the positive class (meaning that the variant is, in fact detected). See **Figure 1** for the pipeline overview. Due to the fact that the task of the model is to differentiate between true and false variants based on alignment, and these features may differ significantly depending on the type of mutation, we decided to train independent models to filter DEL and INS/DUP mutation types.

**Figure 1.**
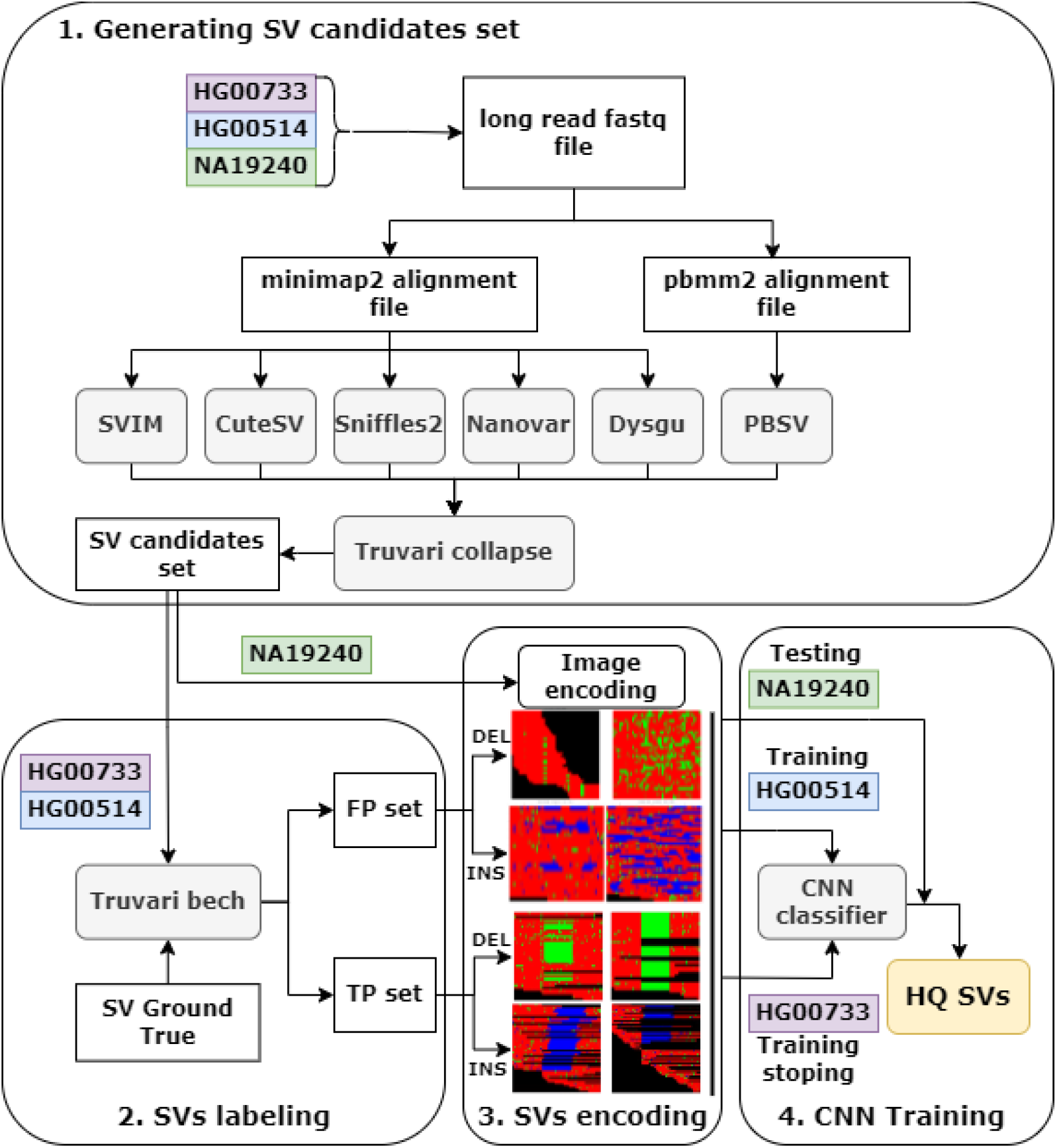
The overview of ConsensuSV-ONT.

### Detection results

In our study, we have used CuteSV ^13^, Sniffles ^14^, Dysgu ^15^, SVIM ^16^, PBSV and Nanovar ^17^ on all the samples, merged their results, and collapsed the merged version to remove redundant variants. Based on the collapsed set, we finally used the ConsensuSV-ONT algorithm. We have decided to compare our work against cnnLSV, and we ran it with a merged set, that was additionally filtered by their script for removing redundant variants (see **Supplementary Materials**). Finally, we compared the detection results of 9 methods with the ground truth set using Truvari. See **Table 1** for the performance of single SV callers, collapsed set, ConsensuSV-ONT, and cnnLSV.

**Table 1.** Detailed detection performance on NA19240, HG00514, HG00733 raw datasets. Bold values represent the best scores.

| Dataset |  | NA19240 |  |  | HG00514 |  |  | HG00733 |  |  |
| --- | --- | --- | --- | --- | --- | --- | --- | --- | --- | --- |
| Type | Method | Prec | Recall | F1 | Prec | Recall | F1 | Prec | Recall | F1 |
| INS | CuteSV | 0.7392 | 0.4569 | 0.5648 | 0.7563 | 0.4307 | 0.5488 | 0.7673 | 0.3989 | 0.5249 |
|  | Dysgu | 0.7262 | 0.4841 | 0.581 | 0.7327 | 0.4618 | 0.5666 | 0.752 | 0.4453 | 0.5594 |
|  | SVIM | 0.7954 | 0.4115 | 0.5424 | 0.8095 | 0.3719 | 0.5097 | 0.8361 | 0.3563 | 0.4997 |
|  | PBSV | 0.6317 | 0.467 | 0.537 | 0.6442 | 0.4413 | 0.5238 | 0.6556 | 0.4301 | 0.5194 |
|  | Nanovar | 0.324 | 0.1995 | 0.2469 | 0.3132 | 0.1921 | 0.2381 | 0.3314 | 0.1861 | 0.2384 |
|  | Sniffles | 0.7665 | 0.326 | 0.4574 | 0.7851 | 0.3015 | 0.4357 | 0.7923 | 0.2929 | 0.4276 |
|  | Collapsed | 0.528 | **0.6585** | 0.5861 | 0.5235 | **0.6329** | 0.573 | 0.5421 | **0.612** | 0.5749 |
|  | ConsensuSV-ONT | 0.7540 | 0.5692 | **0.6487** | 0.7316 | 0.5641 | **0.6371** | 0.7708 | 0.5402 | **0.6352** |
|  | cnnLSV | **0.8101** | 0.3777 | 0.5152 | **0.8326** | 0.3571 | 0.4998 | **0.8513** | 0.3436 | 0.4896 |
| DEL | CuteSV | 0.289 | 0.5738 | 0.3844 | 0.2436 | 0.5776 | 0.3426 | 0.2464 | 0.5578 | 0.3418 |
|  | Dysgu | 0.2846 | 0.6395 | 0.3939 | 0.2382 | 0.646 | 0.3481 | 0.2441 | 0.6306 | 0.352 |
|  | SVIM | 0.3571 | 0.6205 | 0.4533 | 0.306 | 0.6133 | 0.4083 | 0.3191 | 0.6069 | 0.4183 |
|  | PBSV | 0.3849 | 0.6484 | 0.4831 | 0.3169 | 0.6436 | 0.4246 | 0.3269 | 0.636 | 0.4318 |
|  | Nanovar | 0.3116 | 0.4687 | 0.3743 | 0.251 | 0.4854 | 0.3309 | 0.2625 | 0.4664 | 0.3359 |
|  | Sniffles | 0.4971 | 0.497 | 0.4971 | 0.4066 | 0.5011 | 0.4489 | 0.4178 | 0.4909 | 0.4514 |
|  | Collapsed | 0.1711 | **0.7424** | 0.2782 | 0.1365 | **0.7431** | 0.2306 | 0.1431 | **0.7364** | 0.2369 |
|  | ConsensuSV-ONT | **0.6899** | 0.6574 | **0.6733** | **0.6739** | 0.6632 | **0.6685** | **0.6531** | 0.6623 | **0.6577** |
|  | cnnLSV | 0.4921 | 0.5684 | 0.5275 | 0.4248 | 0.5701 | 0.4869 | 0.4337 | 0.5580 | 0.4881 |

The results in the aforementioned table show clearly that ConsensuSV-ONT outperforms each of the single SV callers, as well as the cnnLSV tool. The results obtained by combining the results of single SV callers (collapsed) give the highest recall, but a low F1 score due to low precision, especially for deletions. The superior performance in the case of deletions is seen with precision and F1 score metrics, and in the case of the insertions is seen with F1 score. What’s worth noting is that both ConsensuSV-ONT and cnnLSV are superior to the single tools, supporting our claim even further that consensus approaches are the preferred method for downstream analysis.

### The running time and executive environment

Processing of one sample with mean coverage of ∼30X with 120 Gb RAM and 30 cores by whole pipeline takes around 25 hours. It includes generating alignment, set of SV callings, image encoding and filtering high quality variants with neural network. All this steps are computationally expensive, however our pipeline was developed for processing multiple samples in parallel which could reduce mean processing time for a higher number of samples.

### Comparison of the encoding depending on the sequencing methods

Structural variance detection tools adapted to different sequencing techniques show significant differences in variant prediction ^5^. The differences may be the result of an inaccurate tool that does not detect a given variant or there is noise in a given region that makes it difficult to detect mutations. Visual analysis of images encoded by different sequencing techniques can help understand the origin of differences in mutation detection in the same region using different sequencing techniques. To fully understand the encoding method, we have decided to prepare a small study on how the encoding looks like, depending on the sequencing method used. For this purpose, we selected several mutations from the ground truth set for the HG00514 dataset and used alignment files (bam) to encode regions in the images with our own algorithm. For PacBio sequencing (coverage ∼40X), the bam file (produced by pbmm2) was downloaded directly from the public repository, and for Illumina (coverage ∼74X) and ONT (coverage ∼30X), we downloaded the fastq files and performed alignment using the minimap2 tool. The comparison results are shown in **Figure 2**. Panel A shows a small deletion (53bp) and confirms the occurrence of the mutation across all sequencing techniques. Additionally, we see that PacBio sequencing represents deletions in a highly fragmented manner. This made it impossible for the convolutional model to predict deletions at this location (the values in brackets indicate the value of the probability of mutations occurring at a given position, but only for long-read sequencing). In panel B we can see a large deletion of approximately 10,000 bp, and the encoded image for Illumina sequencing is very different from the others. It does not indicate the deletion in the central part of the image (no green colour), but indicates the lack of data (black colour), which is caused by the lack of reads mapped at least partially along the entire length of the deletion. In the border regions of the beginning and end of the deletion, we see red fragments, which indicate the occurrence of mapping in these regions. For the remaining sequencing types, we note that the mutation was correctly predicted. This suggests that models trained on ONT data may need to be retrained on Illumina data and PacBio data to obtain similar performance in filtering out high-quality variants.

**Figure 2.**
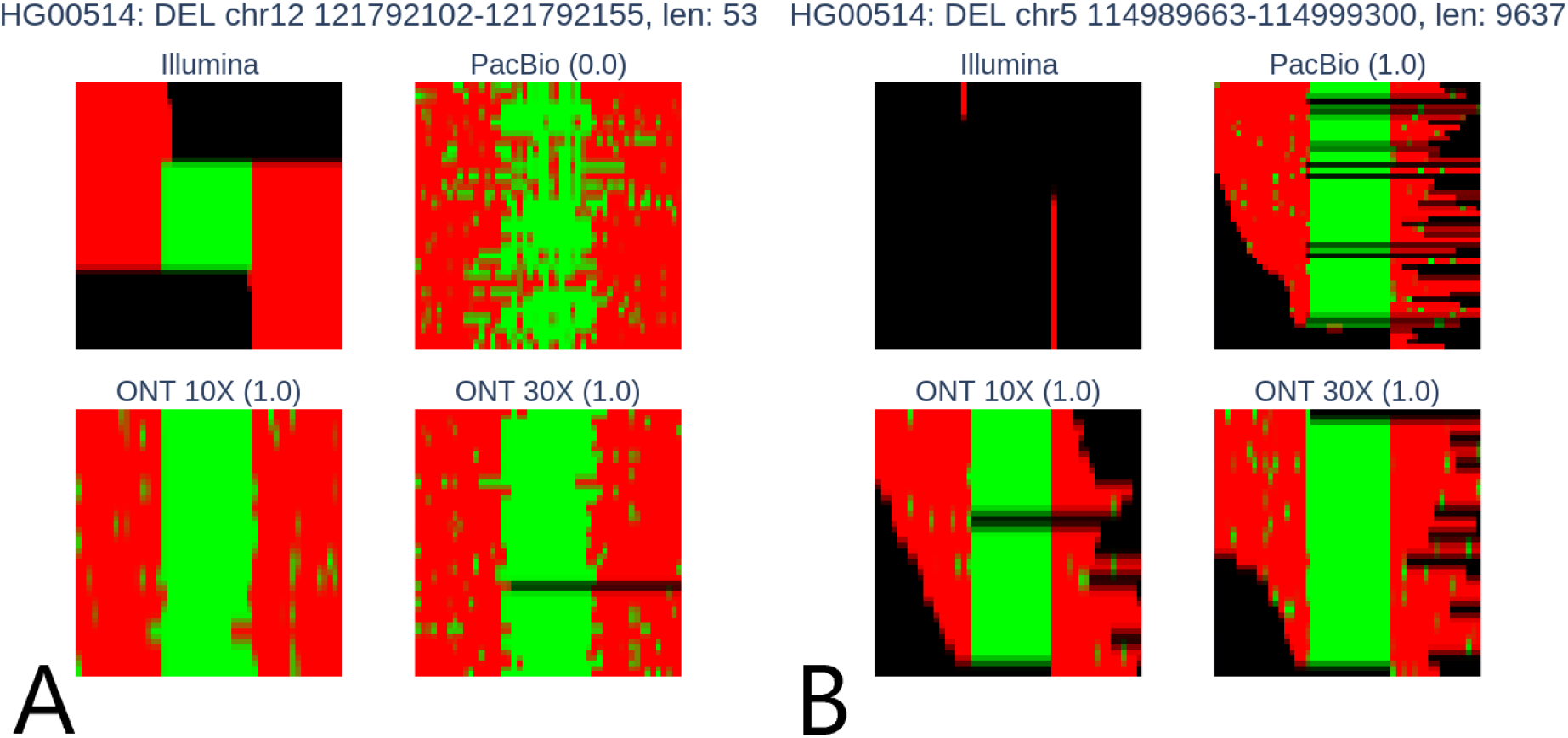
Encoding of example deletions by different sequencing methods. Deletion of chr12 121792102-121792155 could be detected correctly by all sequencing methods (A). Deletion of chr5, 114989663-114999300 cloud be detected correctly only by long-read sequencing methods (B). Values in brackets represent scores predicted by ConsensuSV-ONT, but only for long-read sequencing.

### Analysis of the false positives detected by ConsensuSV-ONT

To fully understand the data we are working on, we have conducted additional analysis of the false positives that our algorithm reports in the HG00514 sample. We have come up with a few possible reasons why the algorithm detects false positives. The first possibility is that a few mutations from the ground truth set were predicted as a single mutation - see **Figure 3** for visualisation.

**Figure 3.**
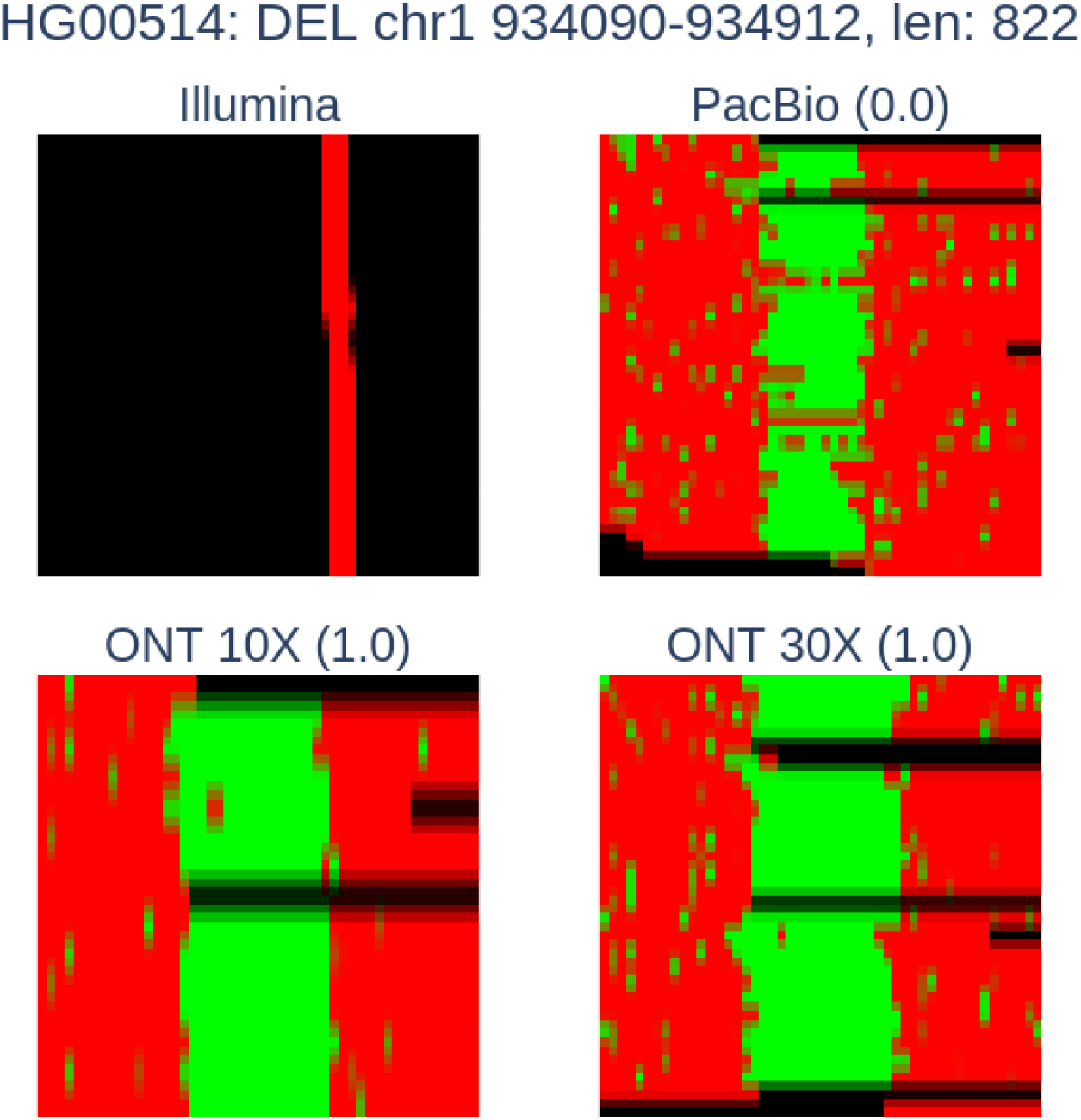
Encoding of merged mutation by different sequencing methods.

Detailed analysis of the region (split into subregions in **Table 2**) showed that the ground truth set reports three separate but very close deletions. The Truvari bench module does not allow interpreting this as the same deletion, and this mutation was classified as a false positive example.

**Table 2.**
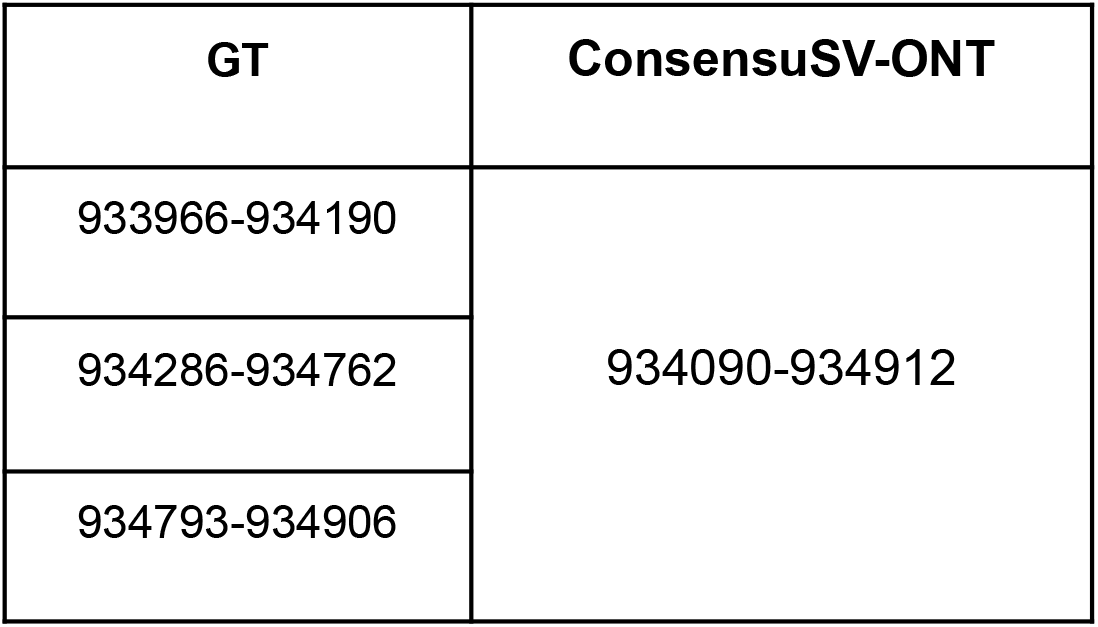
Comparison of mutation detected by ConsensuS-ONT and overlapped mutations from ground truth set.

The second option is that there may have been some overlap, but not enough to consider it the same mutation, as shown in **Figure 4**. Panel A shows deletion from the ground truth set and how it is encoded using sequencing from different methods. We can see that ONT 10X and 30X show shifting deletion to the right side. For PacBio, we can see many scattered short deletion fragments that start at around 1/3 of the length and are present on the entire right side of the image. Panel B shows that ConsensuSV-ONT predicted deletion region which was overlapped with a deletion from Panel A. In the case of ONT, both 10X and 30X, we can see the deletion in the central part of the image. For PacBio, we can see scattered deletion, but it concentrates around the centre of the image. This example suggests the advantage of alignment performed on ONT data over alignment from PacBio data.

**Figure 4.**
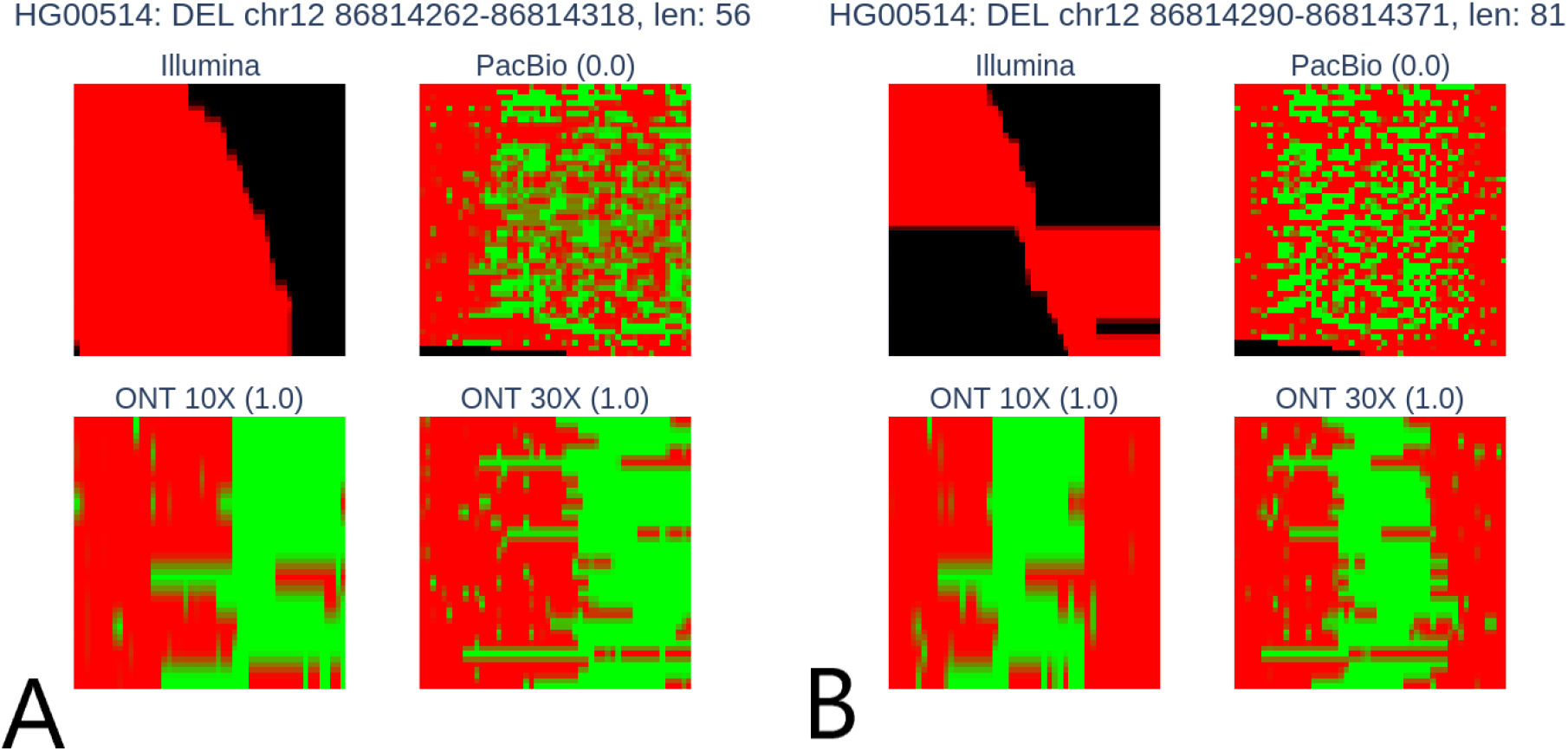
Comparison of the encoding of the deletion reported in the ground truth set, and reported by ConsensuSV-ONT. Ground truth deletion (A) shows the lack of mapping (green color) in the central and right parts of the image. Panel B shows ConsensuSV-ONT reported deletion - and seems to be more stable, as it shows mapping (green colour) in the middle part of the image.

The last example and reason for false positives represent a totally new deletion that was not confirmed by PacBio, Illumina, and 10X ONT. They were not detected because of a lack of reads supporting this deletion in alignment. It is difficult to determine whether the detection of these mutations indicates an advantage or imperfection of the ONT sequence over the others (see **Figure 5**).

**Figure 5.**
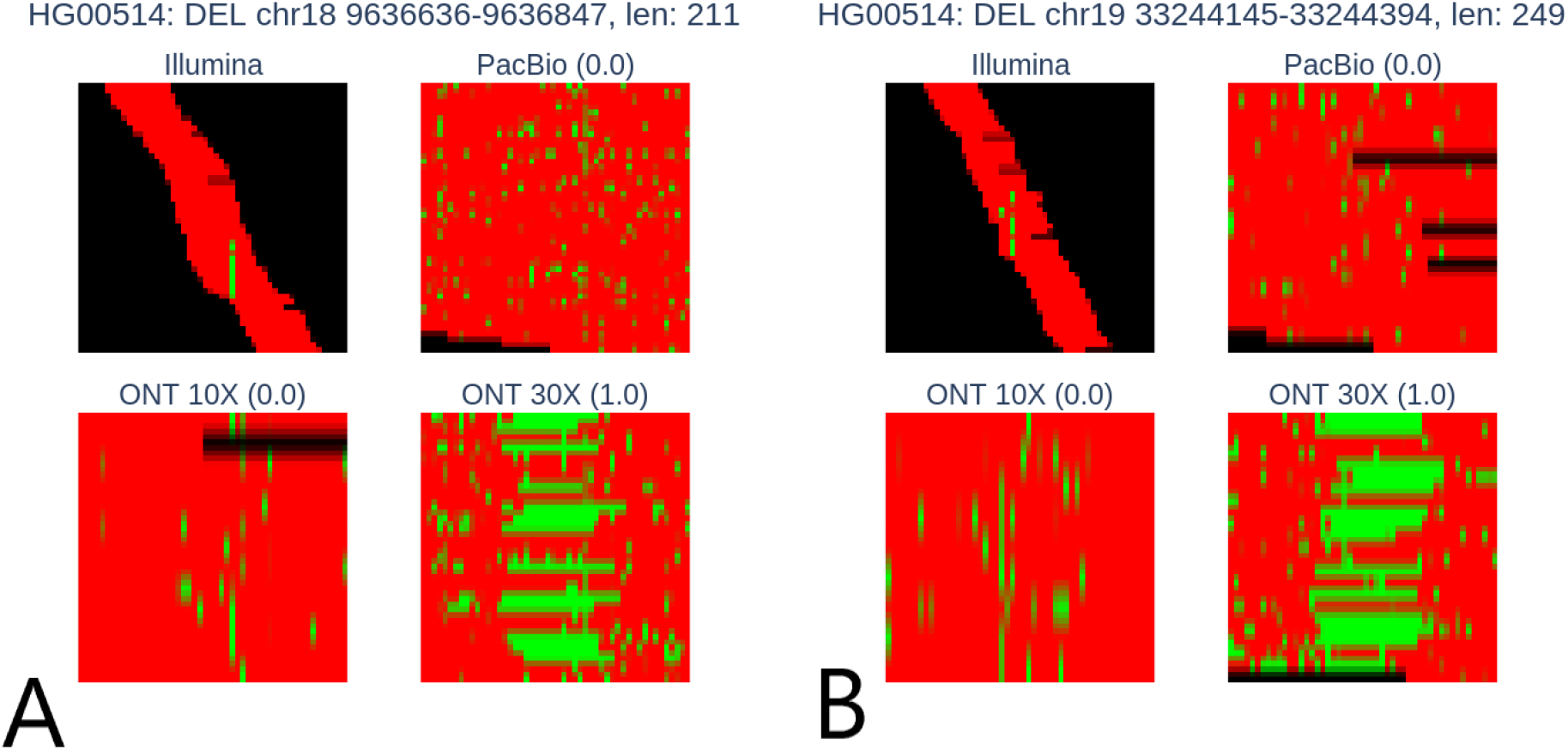
New deletions detected by ConsensuSV-ONT.

## Methods

### Datasets

We obtained raw ONT, publicly available samples (HG00733, HG00514, and NA19240) from the Human Genome Structural Variation Consortium (HGSVC)^5^, aligned them to reference hg38 using minimap2 ^3,18^, and pbmm2 ^6^ and obtained bam files (pbmm2 alignment file is needed as input to the PBSV algorithm). Subsequently we ran Sniffles ^6,14^, CuteSV ^13^, SVIM ^16^, Dysgu ^15^, Nanovar ^15,17^ on the bam generated by minimap2 ^18^ and PBSV on bam file generated by pbmm2 ^6^ to obtain .vcf (Variant Call Format) files with predicted SVs. Detailed information about coverage and average read length for all datasets are in **Table 3**.

**Table 3.** Coverage per sample through datasets.

| Dataset | HG0733 | HG00514 | NA19240 |
| --- | --- | --- | --- |
| Coverage | 30,54 | 31,56 | 31,12 |
| Mean read length | 11,488.5 | 12,512.6 | 15,949.4 |

The minimal SV length was set to 50 (as we consider shorter to be indels). We noticed that with the increase of the ONT sequencing depth, the number of false positive variants increased. Therefore, we decided to pre-filter the variants using metrics provided independently for each variant calling algorithm (see **Supplementary Materials**). Filtered variants were merged and collapsed using Truvari ^12^.

### SV candidates labelling

In order to train a classification model - convolutional neural network (CNN), we needed labelled data. For this purpose, we used Truvari bench software with default parameters, which divided the SV candidate set into two sets: True Positive (TP), i.e. variants from SV candidates that had their counterparts in the ground true set, and False Positives (FP), i.e. variants from the set of SV candidates that had no counterpart in the ground true set. The data marked in this way was used to train a classifier to distinguish real and false structural variants.

### Model training

Based on the fact that we had three ONT samples, we decided to cross-validate our model. This means that we trained 6 deletion and 6 insertion models. In each cross-validation iteration, the training dataset was used to train CNN classifiers. At the end of each epoch, we predict scores for variants from the validation set. Subsequently, we filter out variants with a predicted score higher than the 0.5 threshold and compare this set with ground truth using the Truvari bench module. Based on these validation results, we decide when to stop training to avoid overfitting. We select the model that obtained the best result for the validation set and perform the final tests using the test set - and only those are reported in this work.

### Variants encoding

The tools used to generate SV candidates have various algorithms that, based on alignment, try to predict the positions of mutations in relation to the reference genome. However, there is a very large difference in the results obtained by different tools. Their effectiveness was assessed using different data sets (real and artificial) with different sequencing methods, different depths, and different data preprocessing methods. This makes it impossible to clearly determine the best tool. The idea of the consensus method is based on analyzing all possible indications of variants selected by different algorithms in order to determine common features.

We propose our own approach, similar in concept to cnnLSV, of encoding structural variants into an image based on the alignment information. We decided to encode the images as a matrix with dimensions (50, 50, 3), where 50 is the height and width of the image, and 3 is the number of channels (and is equivalent to RGB space). We decided to use this encoding because of the ease of the visualisation, as well as it would be impossible to encode it within one channel - as there are four important pieces of information we want to preserve in encoding the alignments into the image: mapping, no mapping/deletion and insertion. In the case of two-dimensional coding, e.g. matrix (50, 50), we would have to represent this different information as a continuous value in the range, e.g. 0-1, and it would be impossible to distinguish them.

Encoding the image involves extracting reads of appropriate quality that are mapped to the mutation region and its surroundings (the window is expanded by the size of the mutation on both sides). We decided to do this as there is a possibility of shifting the alignments in relation to the variant. Initially, the image has a size of (SVLEN*3, number of reads, 3), but at the end, it is transformed using the OpenCV library to the previously discussed size (50, 50, 3). The coding also supports the so-called split-reads, i.e. cases where one read is mapped to two distant regions on the same chromosome. This allows the detection of large deletions.

### Model training

The convolutional neural network was implemented as a binary classifier. In order to monitor the effectiveness of the model, we have implemented a special module enabling the calculation of precision, recall and F1 score metrics using the Truvari. After each epoch of training of the neural network, we calculate the metrics using an independent validation set that does not participate in the training process. Thanks to this, we can control whether the model is overfitting, and in such a case, we stop training the model. A chart including the metrics is shown in **Figure 6**. Models were trained using cross-validation to explore the ability to successfully predict one person’s data from a model trained on another person. We can see that after the second epoch, the model gained the ability to generalise well and began to overfit, which meant that the AUC (**Figure 6B**) of the training set kept increasing; however, the metric calculated for the validation set did not improve. Similarly, the loss on the validation set (**Figure 6C**) stopped decreasing after the 3rd epoch. The learning rate was reduced during training due to the lack of significant improvements in metrics during model training.

**Figure 6.**
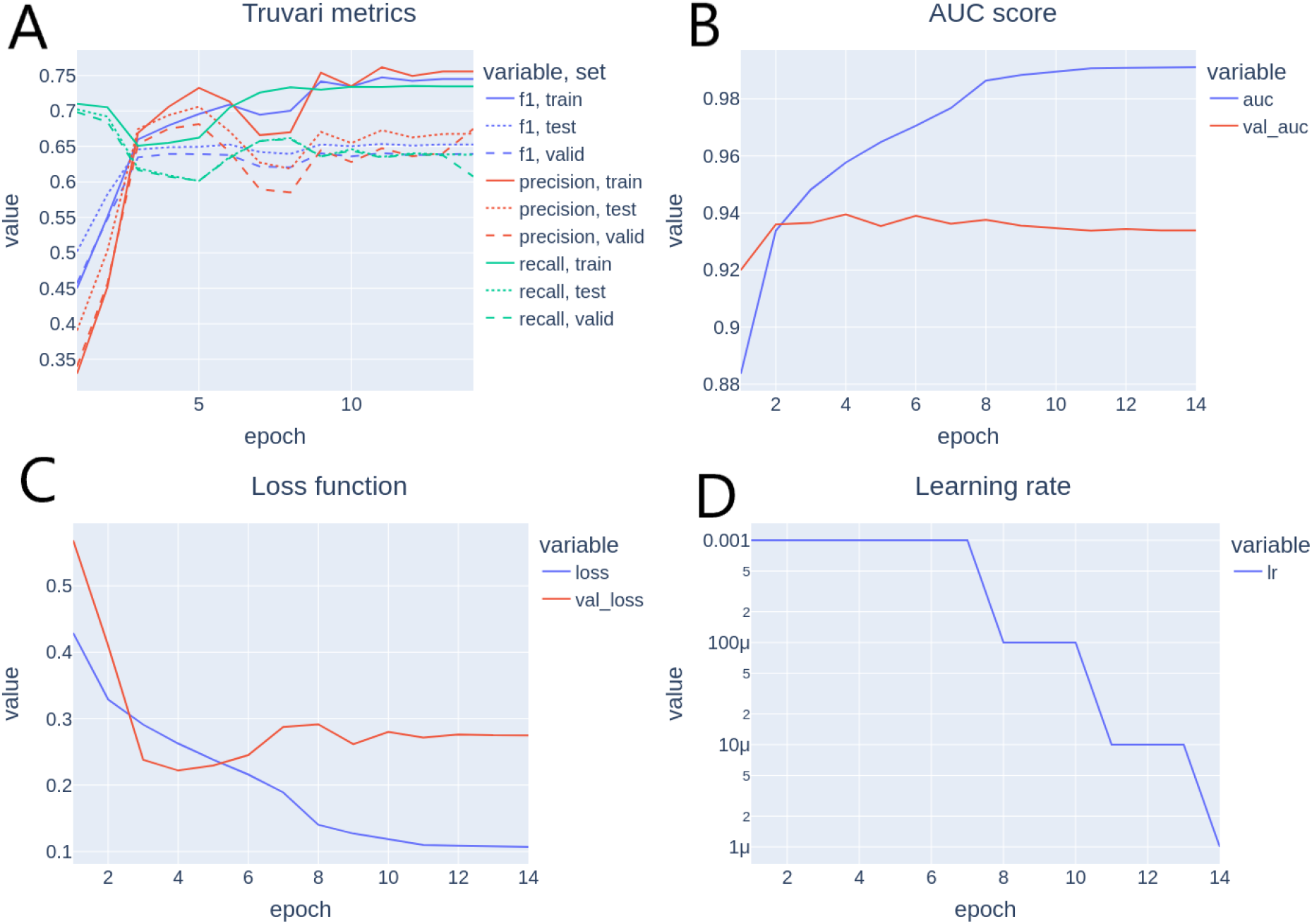
Model train history. Truvari metrics (A) were calculated using Truvari software at the end of each epoch for each dataset. AUC score (B) was calculated only for the training and validation set. Loss function (C) for training and validation set. Learning rate (D) was reduced during training.

### Pipeline

Similarly to the previously published Illumina version of the software ^9^, the implementation is divided into two modules - the first one, ConsensuSV-ONT-core, is used for getting the consensus (by CNN model) out of the already-called SVs, taking as an input vcf files, and returns a high-quality vcf file. The second module is a wrapper for the first one, called ConsensuSV-ONT-pipeline, which is the complete out-of-the-box solution using as the input raw ONT fastq files, and returning as the output the complete lists of high-quality SVs with minimal user involvement. We provide a Docker image with a complete environment, including a nextflow ^19^ pipeline enabling the processing of any number of samples in parallel.

### Truvari

Truvari is a tool that enables many useful operations on files that store structural variance. It enables combining several files into one (merge module). Additionally, it allows combining overlapping variants by the collapse module. However, the greatest benefit comes from the bench module, also used in other works ^10^, which allows you to compare two files with structural variance in order to determine the similarity of the variants. The software returns the following counts for TP, FP and FN and the metrics that are indicated in **Table 4**. TP comp represents the number of variants from the compare file that had an equivalent in the base file. TP-base represents the number of variants from the base file that had their equivalent in the compare file. False Positives (FP) represents variants which were in the compare file but not in the base file. False Negatives (FN) represent variants which were in the base file but not in the compare file.

**Table 4.** Truvari metrics.

| Metric | Formula |
| --- | --- |
| Precision | $\frac{TP\ comp}{(TP\ comp + FP)}$ |
| Recall | $\frac{TP\ base}{(TP\ base + FN)}$ |
| F1 score | $\frac{2 * recall * precision}{recall + precision}$ |

## Discussion

The task of detecting accurate and reliable structural variants is not trivial by any means. In our work, we propose ConsensuSV-ONT tool for long-read SV detection that uses a consensus of independent tools to provide the end user with the best results available. We demonstrated better performance than other tools used for long-read sequencing and created the whole environment with an automated pipeline for sample processing. Our findings suggest that there is a need for a more detailed comparative analysis of mutation detection using different tools and between different sequencing methods. With the expansion of long-read sequencing, our method can be proven to be worthwhile - as it automates most common steps and allows easy detection of the variants. With the progress of multiple population studies on ONT data ^8^, it can bring great benefits.

## Code availability

The code of ConsensuSV-ONT is available at: https://github.com/SFGLab/ConsensuSV-ONT-pipeline/

## Authors’ contributions

A.P. implemented the algorithm under D.P.’s supervision. A.P. performed the analysis with the help of M.C. and S.G. A.P. wrote the manuscript with the help of M.C. under D.P.’s supervision. All authors read and approved the final manuscript.

## Competing Interests

None to declare.

## Funding

This work has been supported by National Science Centre, Poland (2019/35/O/ST6/02484 and 2020/37/B/NZ2/03757). It was co-supported by the Polish National Agency for Academic Exchange (PPN/STA/2021/1/00087/DEC/1). The work has been co-supported by Enhpathy - “Molecular Basis of Human enhanceropathies” funded by the European Union’s Horizon 2020 research and innovation programme under the Marie Sklodowska-Curie grant agreement No 860002 and National Institute of Health USA 4DNucleome grant 1U54DK107967-01 “Nucleome Positioning System for Spatiotemporal Genome Organization and Regulation”. Research was co-funded by the Warsaw University of Technology within the Excellence Initiative: Research University (IDUB) programme. Computations were performed thanks to the Laboratory of Bioinformatics and Computational Genomics, Faculty of Mathematics and Information Science, Warsaw University of Technology, using the Artificial Intelligence HPC platform financed by the Polish Ministry of Science and Higher Education (decision no. 7054/IA/SP/2020 of 2020-08-28).

## Notes

https://github.com/SFGLab/ConsensuSV-ONT-pipeline/tree/main

### Competing Interest Statement

The authors have declared no competing interest.

## References

1. Lander, E. S. et al. Initial sequencing and analysis of the human genome. Nature 409, 860–921 (2001).

2. Sudmant, P. H. et al. An integrated map of structural variation in 2,504 human genomes. Nature 526, 75–81 (2015).

3. Hollox, E. J., Zuccherato, L. W. & Tucci, S. Genome structural variation in human evolution. Trends Genet. 38, 45–58 (2022).

4. Chiliński, M., Sengupta, K. & Plewczynski, D. From DNA human sequence to the chromatin higher order organisation and its biological meaning: using biomolecular interaction networks to understand the influence of structural variation on spatial genome organisation and its functional effect. in Seminars in Cell & Developmental Biology vol. 121 171–185 (Elsevier, 2022).

5. Chaisson, M. J. P. et al. Multi-platform discovery of haplotype-resolved structural variation in human genomes. Nat. Commun. 10, 1784 (2019).

6. Roberts, R. J., Carneiro, M. O. & Schatz, M. C. The advantages of SMRT sequencing. Genome Biol. 14, 405 (2013).

7. Jain, M., Olsen, H. E., Paten, B. & Akeson, M. The Oxford Nanopore MinION: delivery of nanopore sequencing to the genomics community. Genome Biol. 17, 239 (2016).

8. Gustafson, J. A. et al. Nanopore sequencing of 1000 Genomes Project samples to build a comprehensive catalog of human genetic variation. medRxiv 2024.03.05.24303792 (2024) doi:10.1101/2024.03.05.24303792.

9. Chiliński, M. & Plewczynski, D. ConsensuSV-from the whole-genome sequencing data to the complete variant list. Bioinformatics 38, 5440–5442 (2022).

10. Ma, H., Zhong, C., Chen, D., He, H. & Yang, F. cnnLSV: detecting structural variants by encoding long-read alignment information and convolutional neural network. BMC Bioinformatics 24, 119 (2023).

11. Dierckxsens, N., Li, T., Vermeesch, J. R. & Xie, Z. A benchmark of structural variation detection by long reads through a realistic simulated model. Genome Biol. 22, 342 (2021).

12. English, A. C., Menon, V. K., Gibbs, R. A., Metcalf, G. A. & Sedlazeck, F. J. Truvari: refined structural variant comparison preserves allelic diversity. Genome Biol. 23, 271 (2022).

13. Jiang, T. et al. Long-read-based human genomic structural variation detection with cuteSV. Genome Biol. 21, 189 (2020).

14. Smolka, M. et al. Detection of mosaic and population-level structural variants with Sniffles2. Nat. Biotechnol. (2024) doi:10.1038/s41587-023-02024-y.

15. Cleal, K. & Baird, D. M. Dysgu: efficient structural variant calling using short or long reads. Nucleic Acids Res. 50, e53 (2022).

16. Heller, D. & Vingron, M. SVIM: structural variant identification using mapped long reads. Bioinformatics 35, 2907–2915 (2019).

17. Tham, C. Y. et al. NanoVar: accurate characterization of patients’ genomic structural variants using low-depth nanopore sequencing. Genome Biol. 21, 56 (2020).

18. Li, H. Minimap2: pairwise alignment for nucleotide sequences. Bioinformatics 34, 3094–3100 (2018).

19. Di Tommaso, P. et al. Nextflow enables reproducible computational workflows. Nat. Biotechnol. 35, 316–319 (2017).

